# Stoichiometry of HIV-1 Envelope Glycoprotein Protomers with Changes That Stabilize or Destabilize the Pretriggered Conformation

**DOI:** 10.1101/2024.10.25.620268

**Authors:** Zhiqing Zhang, Saumya Anang, Qian Wang, Hanh T. Nguyen, Hung-Ching Chen, Ta-Jung Chiu, Derek Yang, Amos B. Smith, Joseph G. Sodroski

**Affiliations:** Department of Cancer Immunology and Virology, Dana-Farber Cancer Institute, Boston, Massachusetts 02215, USA; Department of Microbiology, Harvard Medical School, Boston, Massachusetts 02115, USA; Department of Chemistry, University of Pennsylvania, Philadelphia, Pennsylvania 19104, USA

**Keywords:** Human immunodeficiency virus, Env, trimer, gp120, gp41, subunit interaction, mutation, cold inactivation, CD4-mimetic compound, State 1

## Abstract

During human immunodeficiency virus (HIV-1) entry into host cells, binding to the receptors, CD4 and CCR5/CXCR4, triggers conformational changes in the metastable envelope glycoprotein (Env) trimer ((gp120-gp41)_3_). CD4 binding induces Env to make transitions from its pretriggered conformation (PTC) to more “open” conformations that are sensitive to inhibition by antibodies, CD4-mimetic compounds (CD4mcs) and exposure to cold. Changes in functional membrane Envs have been identified that either stabilize or destabilize the PTC. Here, we investigate the stoichiometric requirements for the PTC-stabilizing and -destabilizing changes in the Env protomers. To this end, we generated viruses bearing Envs with mixed protomers exhibiting different degrees of PTC stability and determined the sensitivity of the viruses to cold (0°C) and, in some cases, to a CD4mc. The number of stabilized Env protomers required to achieve stabilization of the PTC was inversely related to the degree of PTC stabilization that resulted from the introduced Env change. For strongly stabilizing Env changes, modification of a single protomer was sufficient to achieve PTC stabilization; given adequate stability, the modified protomer can apparently constrain the conformation of the other two protomers to maintain the PTC. Weakly stabilizing Env changes needed to be present in all three protomers to achieve efficient stabilization of the PTC. In many cases, the PTC was disrupted when destabilizing changes were present in only a single protomer. These complementary results suggest that conformational symmetry among the protomers of the functional Env trimer is conducive to the integrity of the PTC.

**IMPORTANCE:** The human immunodeficiency virus (HIV-1) envelope glycoprotein (Env) trimer consists of three protomers. In response to receptor binding, the flexible Env changes its conformation to mediate virus entry into host cells. The shape-shifting ability of Env also contributes to HIV-1’s capacity to evade the host immune system. The pretriggered (State-1) conformation (PTC) of Env is an important target for virus entry inhibitors and host antibodies, but is unstable and therefore incompletely characterized. Changes in Env amino acids that either stabilize or destabilize the PTC have been identified. Here, we define how many Env protomers need to be modified by these changes to achieve stabilization or destabilization of the PTC. These results can guide the placement of changes in the HIV-1 Env protomers to control the movement of the Env trimer from the PTC, allowing better characterization of this elusive conformation and testing of its utility in vaccines.

## INTRODUCTION

The human immunodeficiency virus (HIV-1) envelope glycoprotein (Env) trimer is composed of three protomers, each of which consists of a gp120 exterior Env and a gp41 transmembrane Env (1–4). During HIV-1 entry into host cells, gp120 binds the receptors, CD4 and CCR5/CXCR4, and gp41 mediates the fusion of the viral and target cell membranes (1,2,5–10). The Env trimer is metastable and CD4 binding induces conformational changes from the pretriggered (State-1) conformation (PTC) to more “open” default intermediate (State-2) and full CD4-bound (State-3) conformations (11–13). CCR5/CXCR4 binding to the full CD4-bound state triggers additional conformational changes that lead to the formation of a very stable gp41 six-helix bundle (1,2,14–16). The free energy difference between the metastable PTC and the six-helix bundle is used to drive the fusion of the viral and target cell membranes (1,2,16).

Many HIV-1 biological properties are determined by Env triggerability or reactivity, defined as the propensity of Env to undergo transitions from the PTC to States 2/3 (12,17,18). The metastable pretriggered Env resides in a local energy well and the height of the activation energy barrier separating the PTC and the default intermediate state is inversely related to Env triggerability (12,17,18). Env triggerability governs the virus requirements for levels of CD4 on target cells, sensitivity to soluble CD4 and CD4-mimetic compounds (CD4mcs), and susceptibility to inactivation by prolonged exposure to cold (0°C) (12,17–19). Primary HIV-1 strains exhibit a range of Env triggerabilities (12,17,18). Some natural polymorphisms in Env amino acid residues stabilize or destabilize the PTC (12,17,20–30). In several cases, combinations of these individual amino acids result in additive viral phenotypes (20,24,25,28,29). This additive property has allowed the creation of HIV-1 Env variants that are stabilized in the PTC to a degree beyond that of natural virus strains (25). These extreme examples exhibit very stable Env trimers that are resistant to activation/inactivation by sCD4 and CD4mcs and are less efficient at supporting cell-cell fusion and virus entry (25).

Here, we address the following question: How many Env protomers must be modified by PTC-stabilizing or -destabilizing changes for the functional Env trimer to achieve stabilization or destabilization, respectively, of the PTC? By evaluating the phenotypes of viruses with mixed Env trimers, we find that the number of protomers that need to be modified to achieve PTC stabilization depends upon the degree of PTC stability achieved by the introduced Env change(s). Strongly PTC-stabilizing Env changes need to be present in only a single protomer to stabilize the pretriggered Env trimer, revealing interprotomer cooperativity in regulating PTC stability. On the other hand, weakly PTC-stabilizing changes must be present in all three protomers to achieve stabilization of the PTC. The presence of most destabilizing Env changes in a single protomer was sufficient to disrupt the functional pretriggered Env. These complementary results provide valuable insights into the symmetric nature of the functional pretriggered Env conformation and the interprotomer relationships that govern its maintenance and conversion into downstream conformations.

## RESULTS

### HIV-1_AD8_ Env mutants with alterations in the stability of the PTC

HIV-1 Env triggerability/reactivity is a continuous variable, inversely related to the activation energy required to move Env from its metastable pretriggered conformation (PTC) (12,17,18). We assembled a panel of well-characterized HIV-1_AD8_ Env mutants exhibiting a range of triggerabilities, including those with unusually low triggerability (high PTC stability) and those with high triggerability (low PTC stability) (Table 1) (20,22,24,25). In this set of closely matched HIV-1_AD8_ mutants, virus resistance to cold (0°C) exposure and to CD4mcs has been shown to correlate closely with the degree of PTC stabilization (20,24,25). Cold resistance is a particularly useful surrogate measure of PTC stability for these stoichiometric studies, as the assay avoids the use of an Env ligand whose binding could potentially be affected by the introduced Env amino acid changes (see Fig. 1A for example). The HIV-1_AD8_ Env variants selected for study exhibit a wide range of sensitivities to cold inactivation and inhibition by a CD4mc, BNM-III-170 (Table 1). The pretriggered Env stability index, which correlates directly with PTC stability and inversely with Env triggerability (20,24,25), is the product of the half-life of the viral infectivity at 0°C and the IC_50_ values for BNM-III-170. In Table 1, the PTC-stabilized and PTC-destabilized Env variants are ranked according to their pretriggered Env stability indices, which exhibit >1500-fold variation in this panel. The diversity of this panel of HIV-1_AD8_ Env variants will allow evaluation of the impact of differences in the pretriggered Env stability index on Env protomer stoichiometry.

**FIG 1.**
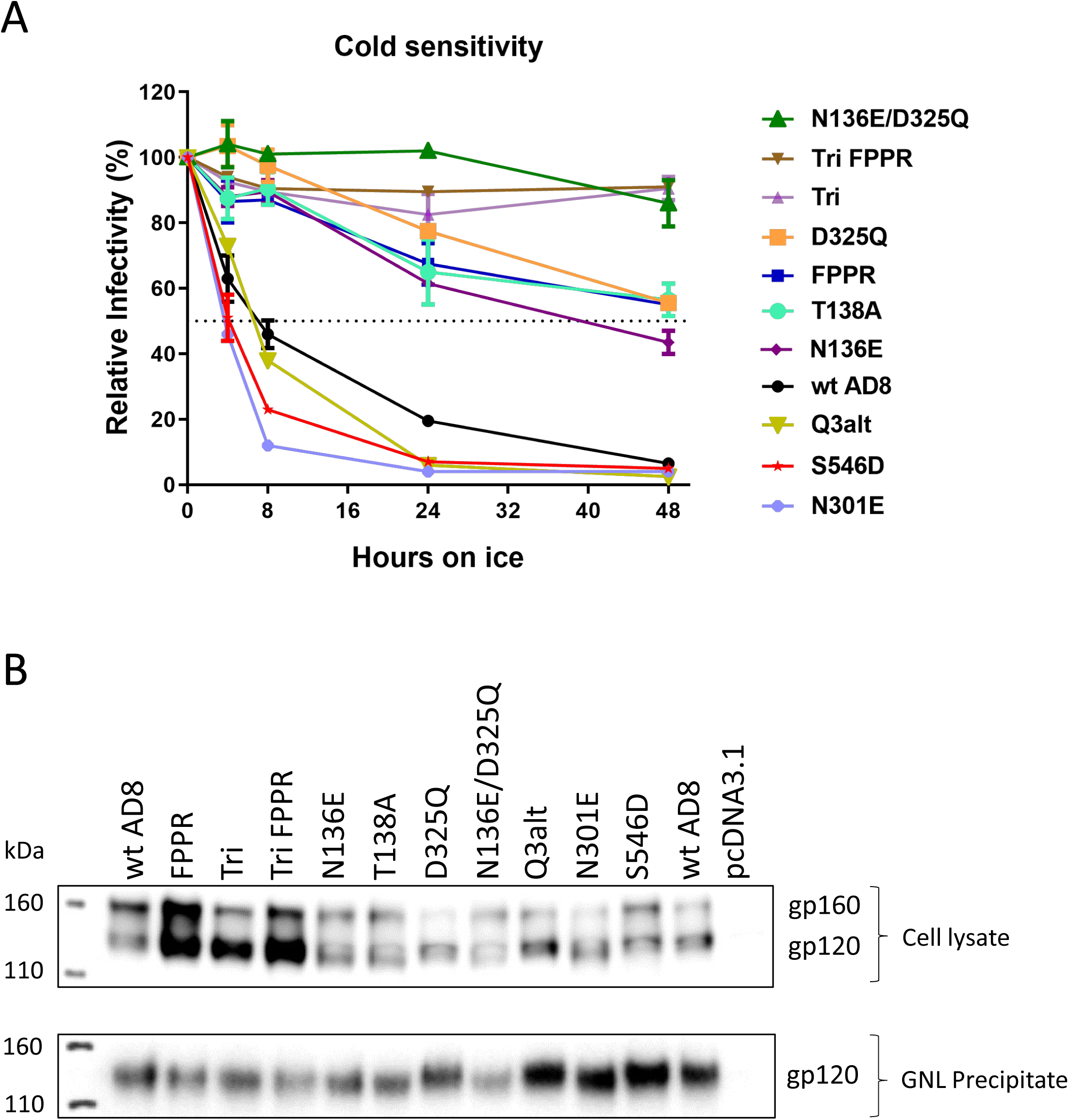
Cold sensitivity, expression and processing of representative Env variants. (A) The cold sensitivity of recombinant HIV-1 pseudotyped with the indicated Env variants is shown. HEK293T cells were transfected with the luciferase-expressing proviral plasmid, pNL4-3.Luc.R-E-, a Tat-expressing plasmid and pSVIIIenv plasmids expressing the indicated wild-type and mutant HIV-1_AD8_ Envs. The pseudoviruses produced in the cell supernatants were incubated on ice (0°C) for the indicated lengths of time and then used to infect TZM-bl cells. After two days, the luciferase activity in the TZM-bl cell lysates was measured. The luciferase activity measured for each virus was normalized to that observed for the same virus sample that was not incubated on ice. The results of a typical experiment are shown, with means and standard deviations derived from triplicate measurements. (B) HOS cells were transfected with plasmids expressing Tat and the wild-type (wt) and mutant HIV-1_AD8_ Envs. Seventy-two hours later, the cells were lysed and the gp120 glycoprotein in the cell medium was captured on Galanthus Nivalis Lectin (GNL)-beads. The clarified cell lysates and GNL precipitates were Western blotted with a polyclonal goat anti-gp120 antibody (Invitrogen).

**TABLE 1.** PTC-stabilized and -destabilized HIV-1AD8 Env variants used in this study.

| HIV-1<sub>AD8</sub> Env<sup>a</sup> | t<sub>1/2</sub> at 0°C (days)<sup>b</sup> | IC<sub>50</sub> BNM-III-170 (μM)<sup>b</sup> | Pretriggered Env stability index<sup>c</sup> | Env processing<sup>d</sup> | Reference |
| --- | --- | --- | --- | --- | --- |
| wild-type | <1 | 4.25 ± 1.47 | 4.25 | +++ |  |
| <u>PTC-stabilized Envs</u> |  |  |  |  |  |
| N136E | <1 | 3.63 ± 2.69 | <3.63 | +++ | 24 |
| Q3 | 1-2 | 2.64 ± 2.20 | 3.96 | +++ | 24 |
| T138A | <1 | 7.29 ± 4.74 | <7.29 | +++ | 24 |
| D325Q | <1 | 8.03 ± 2.32 | <8.03 | +++ | 24,25 |
| Q567K | 1-2 | 6.38 ± 2.69 | 9.57 | +++ | 22 |
| FPPR | 1-2 | 6.87 ± 6.07 | 10.3 | +++ | 24 |
| N136E/D325Q | 1-2 | 10.03 ± 9.19 | 15.0 | +++ | 24,25 |
| A582T | 3-5 | 11.75 ± 4.67 | 47.0 | +++ | 22 |
| Q114E | >5 | >17 | >85 | +++ | 22 |
| FPPR N136E/D325Q | >5 | >21 | >105 | ++ | 24,25 |
| Tri | >5 | >39 | >195 | ++++ | 22 |
| Tri FPPR | >5 | >50 | >250 | ++++ | 25 |
| <u>PTC-destabilized Envs</u> |  |  |  |  |  |
| F317W | <1 | 1.58 ± 0.37 | <1.58 | +++ | 30 |
| Q3alt | <1 | 1.10 ± 0.17 | <1.10 | +++ | 24 |
| Q3-N332T | <1 | 0.99 ± 0.50 | <0.99 | ++++ | 24 |
| N332T | <1 | 0.48 ± 0.39 | <0.48 | ++++ | 24 |
| N301E | <1 | 0.25 ± 0.08 | <0.25 | +++ | 24 |
| S546D | <1 | 0.16 ± 0.07 | <0.16 | +++ | 22 |

<sup>a</sup>The HIV-1<sub>AD8</sub> Env variants used in this study are listed, with the Env residues numbered according to current convention (44).
<sup>b</sup>To ensure an accurate ranking of these mutants, the half-lives of pseudovirus infectivity on ice and the IC<sub>50</sub> values of the CD4mc BNM-III-170 were determined in side-by-side assays, as described in Materials and Methods.
<sup>c</sup>The pretriggered Env stability index is the product of the virus half-life at 0°C and the BNM-III-170 IC<sub>50</sub>. Where a range of cold half-lives is reported, an average value was used to calculate the pretriggered stability index. The PTC-stabilized and PTC-destabilized HIV-1<sub>AD8</sub> Env variants are ranked according to their degree of PTC stabilization or destabilization, respectively.
<sup>d</sup>Env processing was evaluated in HOS cells, as described in Materials and Methods.

Most of the Env mutants selected for this study were expressed and processed at least as well as the wild-type HIV-1_AD8_ Env (Table 1 and Fig. 1B). Some of the Envs (Tri, Tri FPPR, N332T and Q3-N332T) were processed more efficiently than the wild-type HIV-1_AD8_ Env. Only one mutant (FPPR N136E/D325Q) was processed less efficiently than the wild-type HIV-1_AD8_ Env. The potential impact of differences in Env processing will be considered in the interpretation of the results (see below).

### Generation of recombinant HIV-1 with mixed Env trimers

We generated recombinant single-round viruses expressing luciferase that were pseudotyped with mixed Env trimers. The Envs were composed of the wild-type HIV-1_AD8_ Env mixed with either PTC-stabilized or PTC-destabilized mutant Envs in varying proportions. We predict that the phenotype (P) of viruses with mixed Envs, relative to the phenotypes of viruses with 100% wild-type or 100% mutant Envs, is determined by the percentage of Env trimers with a stabilized (or destabilized) PTC in the mixture. P is a function of the fraction (f) of mutant Envs in the mixture and the number of mutant Env protomers (N_p_) required to achieve the PTC-stabilized or -destabilized phenotype (Fig. 2). We assume: 1) a random mixing of Envs according to a binomial distribution; and 2) all-or-none Env stabilization/destabilization. Given these assumptions, the ideal curves on a graph of P versus f are shown in Fig. 2. Fitting the empirical plot of the viral phenotype versus f to these ideal curves, which qualitatively differ for N_p_=1, 2 and 3, allows us to deduce the value of N_p_.

**FIG 2.**
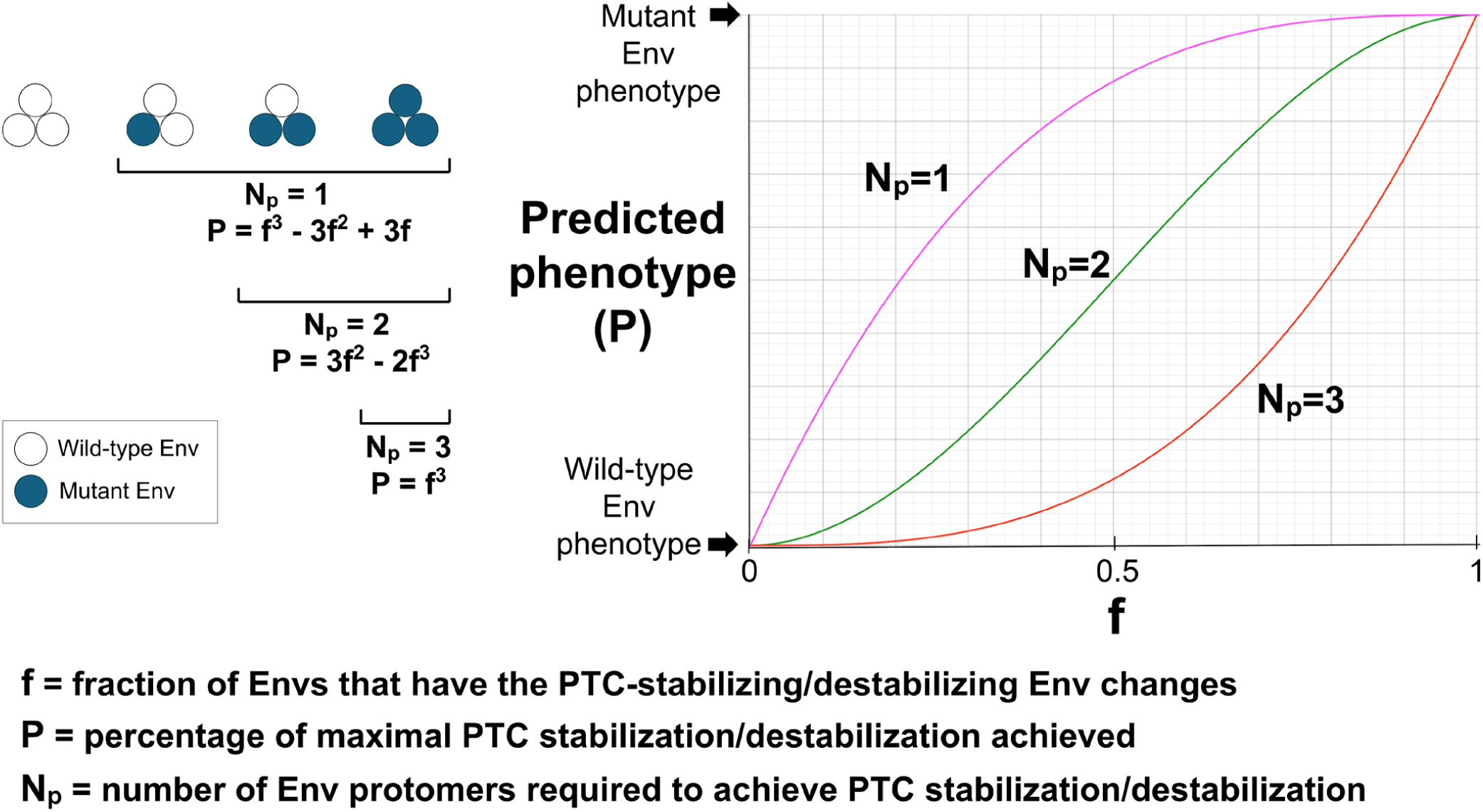
Theoretical analysis of Env protomer stoichiometry for PTC stabilization and destabilization. The left panel considers the random mixing of wild-type Envs (open circles) and PTC-stabilized/destabilized mutant Envs (filled circles) into trimers according to a binomial distribution. The percentage of the Env trimers in the population that determines the PTC-stabilized/destabilized viral phenotype (P) depends on the number of Env protomers (N_p_) required to achieve PTC stabilization or destabilization and on the fraction of mutant Envs (f). We assume that the percentage of the maximal phenotype (defined by the phenotype of viruses with 100% mutant Envs) achieved by the mixed-Env viruses is proportionate to the percentage of PTC-stabilized/destabilized Env trimers in the virus population. The right panel shows the theoretical graphs of P as a function of f for the different N_p_ values. Note that the shape of the f-P curve is distinct for each value of N_p_.

### Protomeric stoichiometry of Envs with PTC-stabilizing changes

Recombinant viruses with mixtures of the wild-type HIV-1_AD8_ Env and PTC-stabilized variant Envs were generated. The infectivity of these viruses was measured after incubation on ice (0°C) for a period of time that resulted in maximal differences between the infectivities of the viruses with 100% wild-type HIV-1_AD8_ Env and 100% mutant Env. Plots of P (in this case, residual virus infectivity after cold exposure) as a function of f (the fraction of mutant Envs) revealed three patterns (Fig. 3). Convex upward f-P curves were observed for three Env variants, Q114E, Tri and Tri FPPR. The Tri and Tri FPPR Envs have multiple PTC-stabilizing changes, one of which is Q114E (20,24,25). The shapes of these f-P curves is consistent with N_p_=1, indicating that the presence of these PTC-stabilizing changes in a single protomer is sufficient to achieve cold resistance. For three of the Env variants (FPPR-N136E/D325Q, A582T and FPPR), the f-P curves were consistent with N_p_=2, suggesting that the presence of these PTC-stabilizing changes in two protomers was sufficient to confer cold resistance. A third pattern of concave upward f-P curves was observed for half (6 out of 12) of the PTC-stabilizing changes.

**FIG 3.**
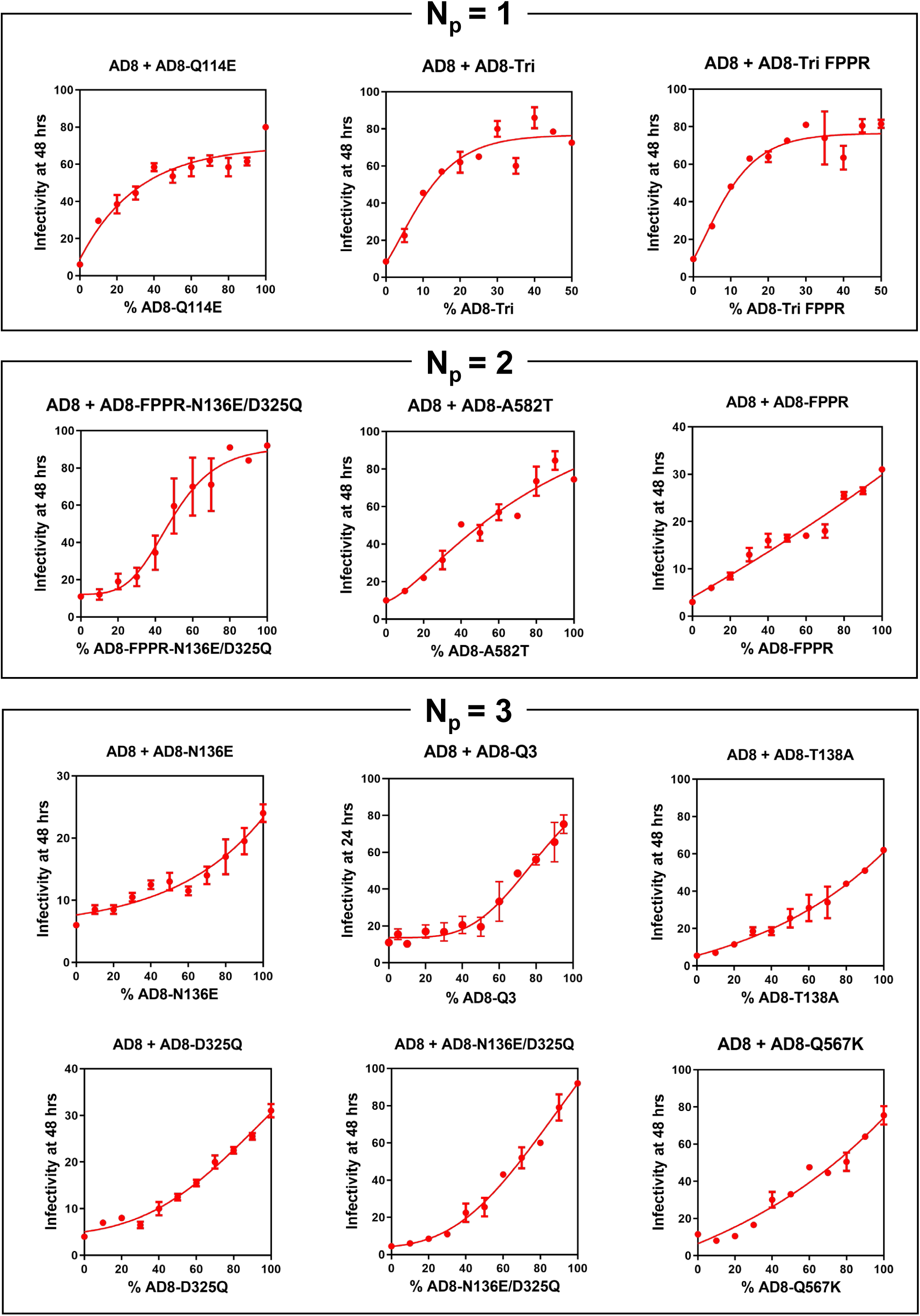
Cold sensitivity of viruses with mixed wild-type and PTC-stabilized Envs. HEK293T cells were transfected with the luciferase-expressing proviral plasmid, pNL4-3.Luc.R-E-, a Tat-expressing plasmid and pSVIIIenv plasmids expressing the wild-type and mutant HIV-1_AD8_ Envs in the indicated proportions. The pseudoviruses produced in the cell supernatants were incubated on ice (0°C) for the indicated times and then used to infect TZM-bl cells. Two days later, the luciferase activity in the TZM-bl cell lysates was measured. The luciferase activity measured for each virus was normalized to the luciferase activity observed for the same virus preparation that had not been incubated on ice. The means and standard deviations derived from triplicate measurements within a typical experiment are shown. The PTC-stabilized Env variants are grouped according to the N_p_ values deduced from the shape of the f-P curves.

These f-P curves indicate that the PTC-stabilizing changes need to be present in all three Env protomers to achieve cold resistance (N_p_=3). These results indicate that the number of Env protomers that need to be modified to achieve cold stability of the virus differ for various PTC-stabilizing Env changes.

High levels of expression and processing of mutant Envs relative to those of wild-type Env could hypothetically lower our estimate of N_p_. As some PTC-stabilized Env mutants like Tri and Tri FPPR exhibit relatively efficient processing and low estimated N_p_ values (20,25) (Fig. 1B, Fig. 3 and Table 1), we wished to evaluate whether mixing these mutants with wild-type HIV-1_AD8_ Env might have biased the outcome. We examined the Envs in pseudoviruses produced by titrating increasing amounts of Tri and Tri FPPR Envs into the wild-type HIV-1_AD8_ Env (Fig. 4A). The levels and processing of the mixed Envs with increasing proportions of Tri and Tri FPPR Envs were similar, particularly when these values were normalized to the levels of Gag p24 capsid protein in the virions. These observations argue against systematic alterations of virion Env levels or processing as a result of mixing the Tri and Tri FPPR Envs with the wild-type HIV-1_AD8_ Env.

**FIG 4.**
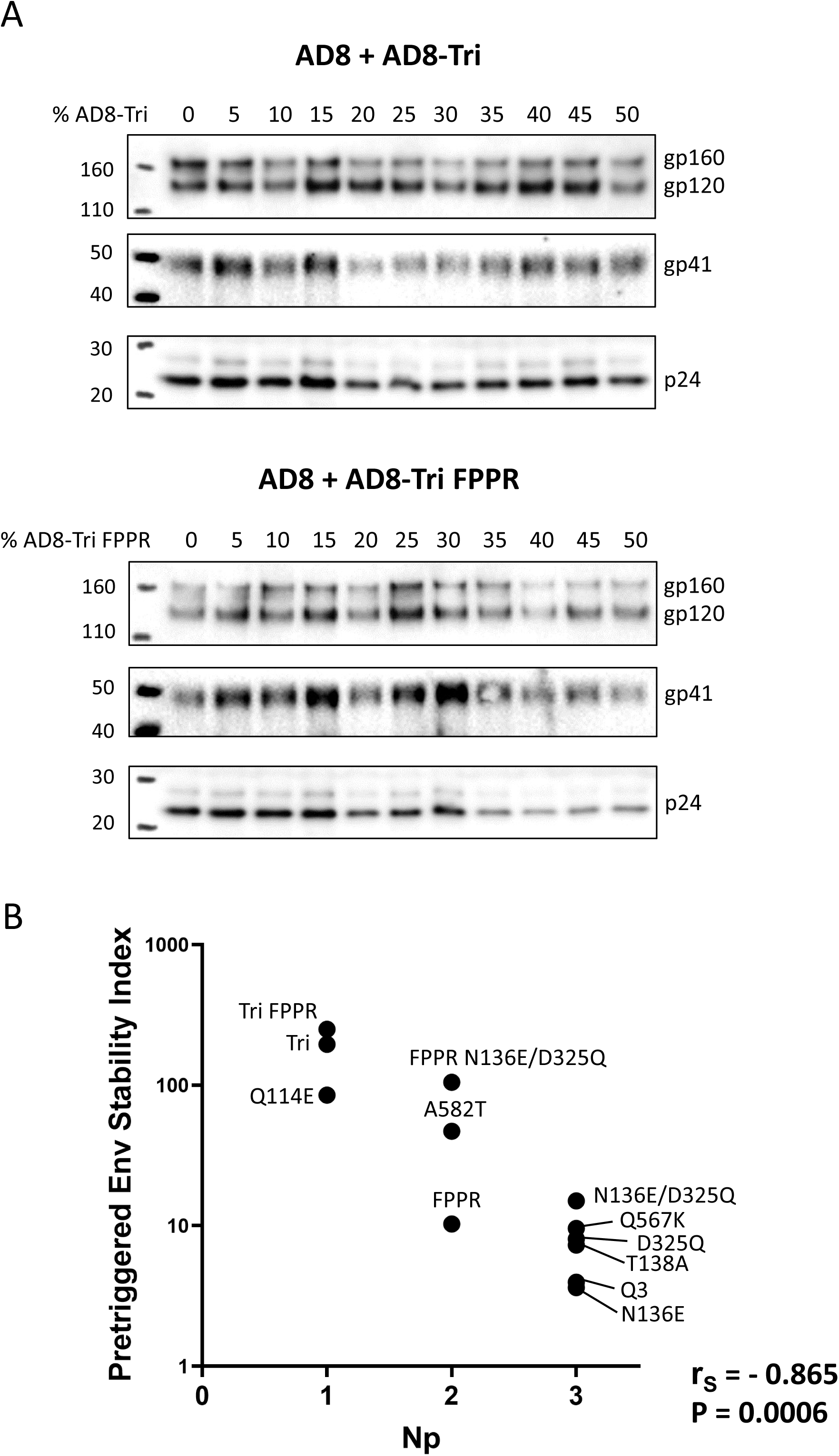
Characterization of viruses with mixed wild-type and PTC-stabilized Envs. (A) Pseudoviruses with the indicated proportions of wild-type and mutant HIV-1_AD8_ Envs were prepared as described in the Figure 3 legend. Clarified cell supernatants were centrifuged at 14,000 x g to pellet the virus particles. Equal volumes of resuspended virus particles were Western blotted with a polyclonal goat anti-gp120 antibody (upper panels), the 4E10 anti-gp41 antibody (middle panels), and a polyclonal rabbit antibody against Gag p55/p24/p17 (lower panels). The results of a typical experiment are shown. (B) The inverse relationship between the N_p_ values deduced from the shapes of the f-P curves in Figure 3 and the pretriggered Env stability indices (Table 1) is shown. The Spearman rank correlation coefficient (r_S_) and two-tailed P value are shown.

Notably, for our panel of PTC-stabilizing Env changes, the N_p_ values deduced from the analysis of the f-P curves inversely correlated with the pretriggered Env stability indices (Spearman r_S_ = −0.865, two-tailed P = 0.0006) (Fig. 4B). Thus, the number of Env protomers that need to be modified to achieve stabilization of the PTC is inversely related to the degree of PTC stabilization that results from the Env change.

Weakly and moderately PTC-stabilizing changes need to be present in multiple protomers (N_p_=3 or 2, respectively) to achieve cold resistance. Strongly PTC-stabilizing Env changes efficiently stabilize the functional PTC against the effects of cold exposure when present in a single protomer.

### Protomeric stoichiometry of Envs with PTC-destabilizing changes

Recombinant viruses with mixtures of wild-type HIV-1_AD8_ Env and PTC-destabilized Env mutants were generated and tested for cold sensitivity. For most of the Env mutants, the presence of the destabilizing changes in a single protomer was sufficient to destabilize the PTC and render the viruses sensitive to incubation at 0°C (Fig. 5). A protomer stoichiometry of N_p_=1 was deduced from the cold sensitivity plots of the Q3alt, N301E, F317W and N332T Env mutants. Of interest, three of these PTC-destabilizing changes (Q3alt, N301E and N332T) involve the loss or shift of an N-linked glycosylation site in gp120 (24). The F317W change alters the tip of the gp120 V3 loop (30).

**FIG 5.**
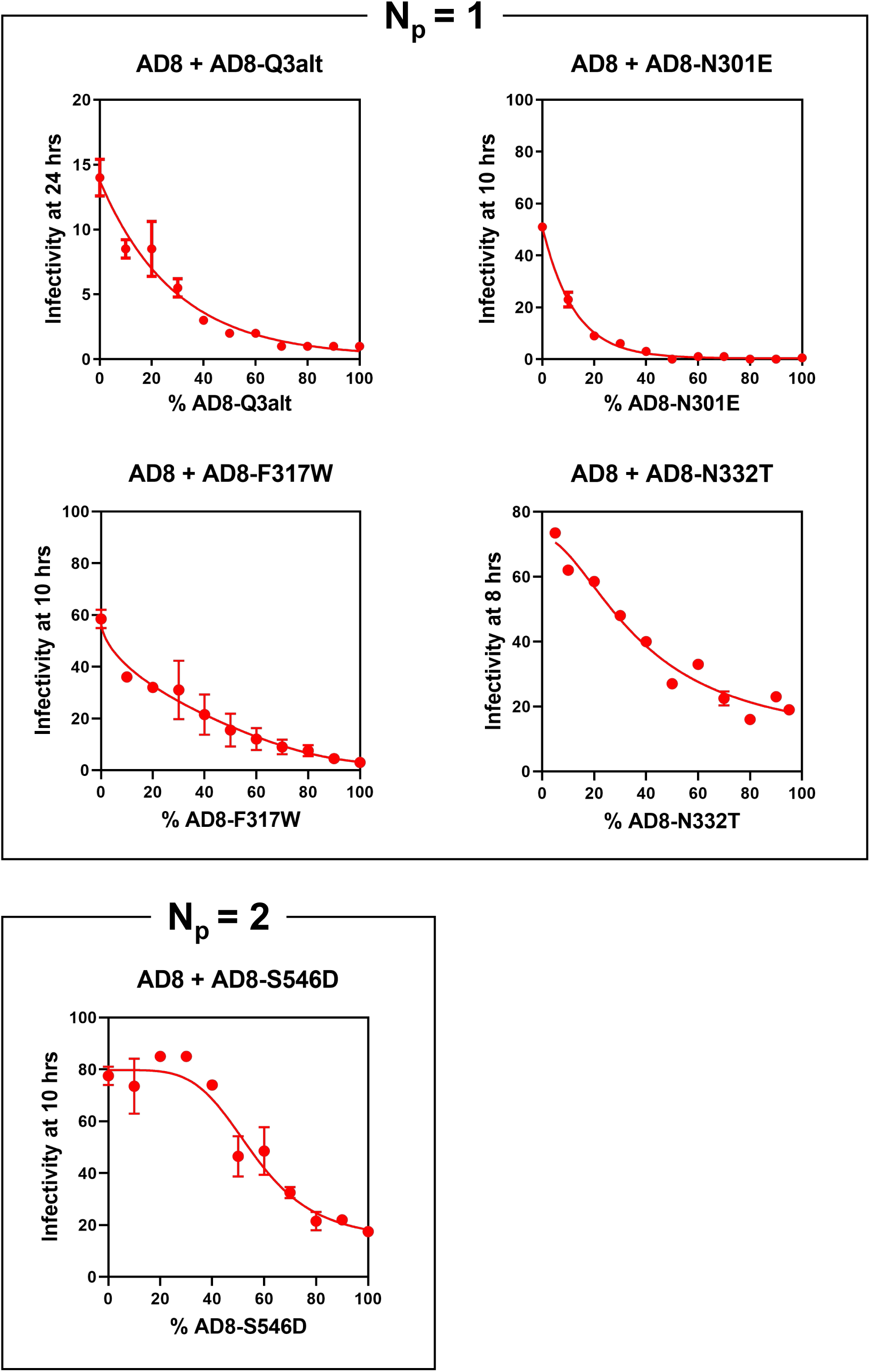
Cold sensitivity of viruses with mixed wild-type and PTC-destabilized Envs. The cold sensitivity of pseudoviruses with mixed wild-type and mutant HIV-1_AD8_ Envs was evaluated as described in the Figure 3 legend. The means and standard deviations derived from triplicate measurements within a typical experiment are shown. The experiments were repeated with comparable results. The PTC-destabilized Env variants are grouped according to the N_p_ values deduced from the shape of the f-P curves.

Another PTC-destabilizing change, S546D, needed to be present in two Env protomers to destabilize the Env trimer and render it sensitive to cold (Fig. 5). Serine 546 is located in the gp41 heptad repeat (HR1_N_) region and the S546D change does not alter a potential N-linked glycosylation site. Thus, the S546D change differs from the other destabilizing Env mutants tested in subunit location in the Env trimer, protomeric stoichiometry and, in some cases, effect on glycosylation.

To evaluate potential relationships between the degree of PTC destabilization and the protomeric stoichiometry (N_p_) of the S546D and the other tested Env variants, we compared the sensitivity of viruses with these Envs to cold, CD4mcs, sCD4-Ig and antibodies. The results are shown in Tables 1 and 2, where the PTC-destabilized Env mutants are arranged in order of decreasing pretriggered Env stability indices. The sensitivity of viruses with these Envs to sCD4-Ig exhibits the same rank order (Table 2); PTC destabilization is expected to increase Env triggerability and inactivation by sCD4-Ig (12,17,19,23–25). The F317W virus exhibited a two-fold increase in sensitivity to sCD4-Ig, but was neutralized by the antibodies tested comparably to the wild-type HIV-1_AD8_ virus. Viruses with the other PTC-destabilized Envs (N332T, N301E and S546D) were more sensitive than the wild-type virus to CD4BS bNAbs, the 10E8.v4 MPER bNAb and the 447-52D anti-V3 pNAb. The S546D virus was not only more sensitive to these antibodies than the N332T and N301E viruses, but also was neutralized by the 19b anti-V3 and 17b CD4i pNAbs. Thus, compared with the other PTC-destabilizing Env changes, the S546D change results in the highest degree of PTC destabilization, based on the viral pretriggered Env stability indices and sensitivities to antibody/sCD4-Ig neutralization. In this set of PTC-destabilized Env variants, greater PTC destabilization is associated with a higher N_p_ requirement.

**TABLE 2.**
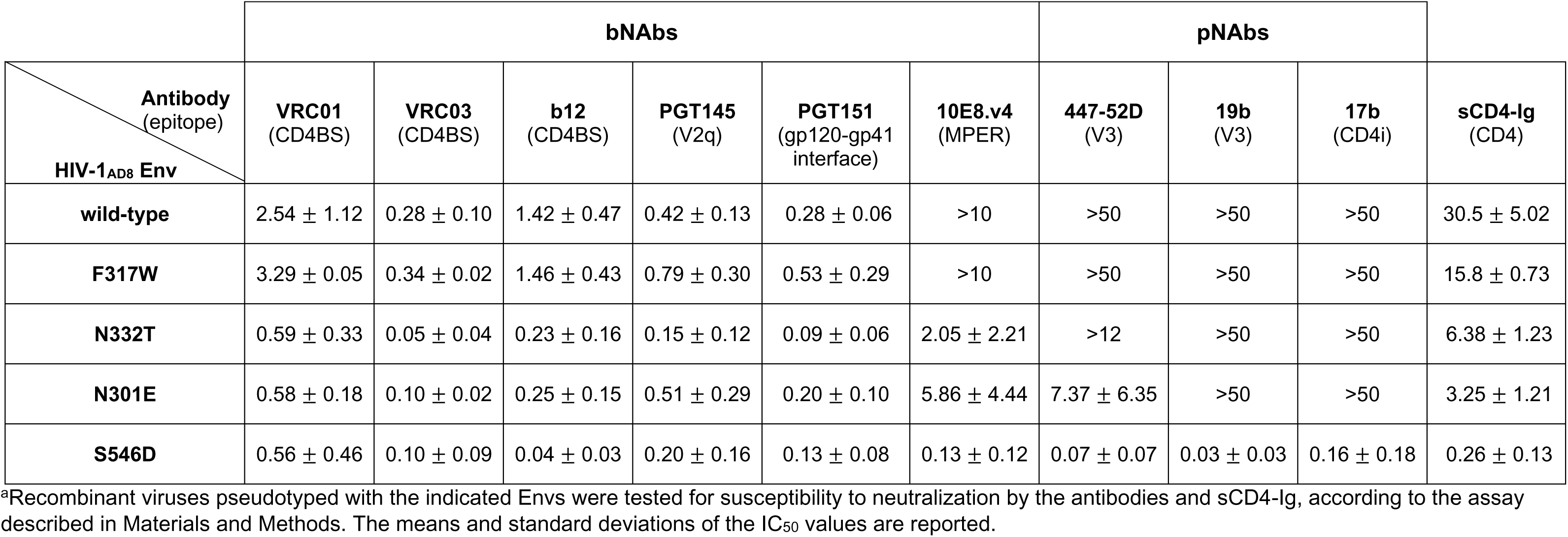
Sensitivity of viruses pseudotyped with PTC-destabilized HIV-1_AD8_ Envs to neutralization by antibodies and sCD4-Ig^a^.

The deduced protomer stoichiometries (N_p_ values) and the relationship of the observed phenotypes to the degree of PTC stabilization/destabilization for the panel of studied Env variants is summarized in Figure 6.

**FIG 6.**
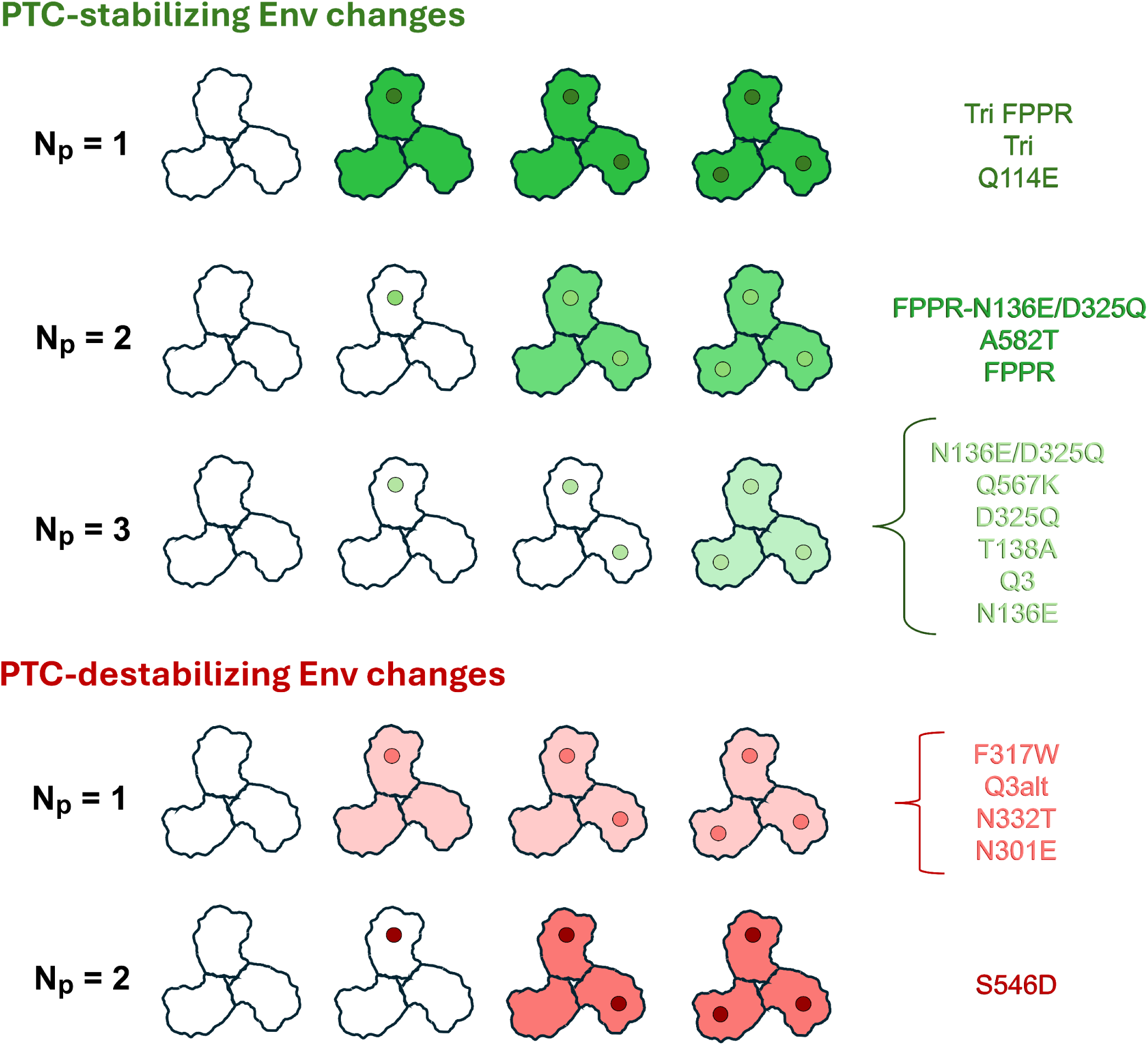
Summary of deduced protomer stoichiometries for PTC-stabilizing and PTC-destabilizing Env changes. A schematic diagram of the HIV-1 Env trimer is shown with PTC-stabilizing changes (green dots) and PTC-destabilizing changes (red dots) in the protomers. The dots are shaded according to the degree of PTC stabilization or destabilization achieved when the respective PTC-stabilizing or PTC-destabilizing Env change is present in all three protomers; the pretriggered Env stability index provides an indication of the degree of PTC stabilization or destabilization achieved by the introduced changes. The specific Env changes assigned to each N_p_ category are listed on the right. Because of the strong relationship between the deduced N_p_ values and the pretriggered Env stability indices (Fig. 4A), the Env changes are roughly ranked from highest to lowest pretriggered Env stability index. The deduced phenotypes of the functional Env trimers are indicated by shading, with cold-resistant trimers in green and cold-sensitive trimers in red. The intensity of the Env trimer shading relates to the degree of PTC stabilization or destabilization.

### Combination of PTC-stabilizing and -destabilizing changes in Env trimers

We wished to examine the effect of PTC-destabilizing Env changes in the context of an Env trimer that has PTC-stabilizing changes in some or all of its protomers. Several variations were tried:

**a) Q3 + Q3-N332T** – In the background of the wild-type HIV-1_AD8_ Env, the N332T change in one protomer (N_p_=1) is sufficient for destabilization of the PTC and increased cold sensitivity (Fig. 5). The magnitude of the cold sensitivity phenotype resulting from the N332T change was reduced when the N332T change was introduced into an Env where all three protomers had the PTC-stabilizing Q3 change (Fig. 7A). Nonetheless, the shape of the f-P curve indicated that the protomer stoichiometry remained at N_p_=1.
**b) Tri FPPR + AD8-N301E** – In the wild-type HIV-1_AD8_ background, the N301E change destabilizes the PTC when present in a single protomer (Fig. 5). Likewise, when the AD8-N301E Env was mixed with the Tri FPPR Env, one protomer of AD8-N301E was sufficient to destabilize the PTC of the Env trimer (Fig. 7A).
**c) Tri FPPR + AD8-Q3alt** – In the wild-type HIV-1_AD8_ background, the Q3alt change in a single protomer is sufficient to destabilize the PTC (Fig. 5). This was also the case when the AD8-Q3alt Env was mixed with the Tri FPPR Env (Fig. 7A).
**d) Tri FPPR + AD8-S546D** – In the wild-type HIV-1_AD8_ background, two S546D protomers are required to destabilize the PTC of Env (Fig. 5). However, when the AD8-S546D Env was mixed with the Tri FPPR Env, one AD8-S546D protomer was apparently able to bring about PTC destabilization (Fig. 7A).

**FIG 7.**
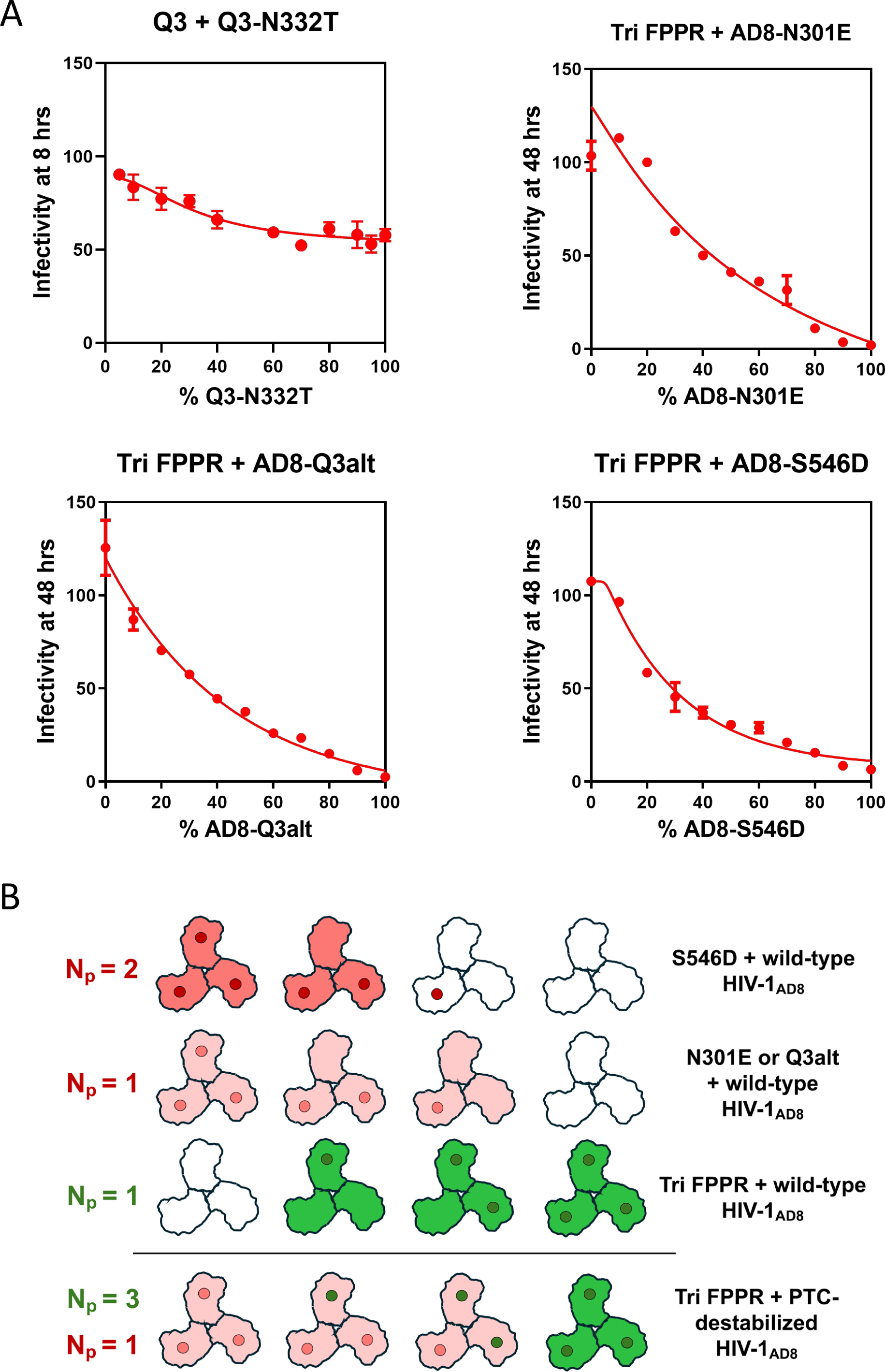
Cold sensitivity of viruses with mixed PTC-stabilized and PTC-destabilized Envs. (A) The cold sensitivity of pseudoviruses with mixed PTC-stabilized and PTC-destabilized Envs was evaluated as described in the Figure 3 legend. The means and standard deviations derived from triplicate measurements within a typical experiment are shown. The experiments were repeated with comparable results. (B) A schematic diagram of the HIV-1 Env trimer is shown with the PTC-stabilizing Tri FPPR changes (green dots) and the PTC-destabilizing (N301E, Q3alt and S546D) changes (red dots) in the protomers. The Env variants in the mixed virus Envs are shown on the right. The viral Env mixtures above the horizontal line include the wild-type HIV-1_AD8_ Env as a partner and thus correspond to those depicted in Figure 6. The deduced N_p_ values of the Tri FPPR Env (green) and the PTC-destabilized Envs (red) are indicated on the left. The deduced phenotypes of the functional Env trimers are indicated by shading, with cold-resistant trimers in green and cold-sensitive trimers in red. Wild-type HIV-1_AD8_ cold sensitivity is indicated by white/unshaded trimers. Note that compared to the mixtures with wild-type HIV-1_AD8_ Env, when the Envs with the PTC-stabilized Tri FPPR changes are mixed with the PTC-destabilized Envs, N_p_=1 for the PTC-destabilized Envs and the N_p_ value of Tri FPPR increases to 3.

**FIG 8.**
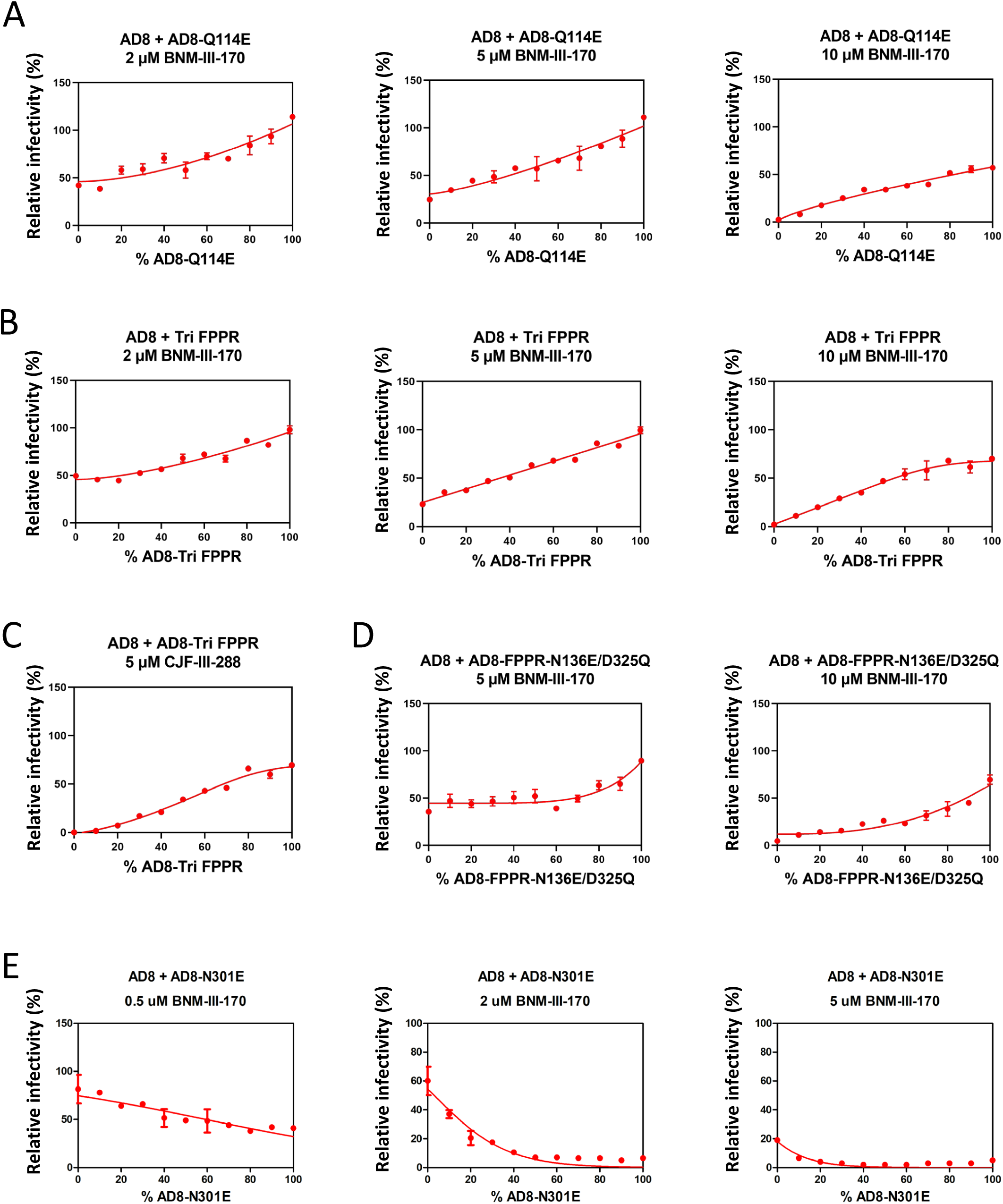
CD4mc inhibition of viruses with PTC-stabilized (or PTC-destabilized) Envs mixed with the wild-type HIV-1_AD8_ Env. Pseudoviruses with mixed wild-type and mutant HIV-1_AD8_ Envs were prepared as described in the Figure 3 legend. The viruses were incubated with the indicated concentrations of the CD4mc BNM-III-170 (A, B, D and E) or the more potent CD4mc CJF-III-288 (C) for 1 hour at 37°C. The virus-CD4mc mixture was then incubated with TZM-bl cells for two days, after which luciferase activity in the target cells was measured. The measured luciferase activity was normalized to that observed for the same virus preparation that had not been treated with the CD4mc. Multiple concentrations of the CD4mc were tested to identify concentration(s) at which the differences between the wild-type and mutant Envs would permit conclusions about the Env protomer stoichiometry. In A, B, D and E, the results obtained with different concentrations of BNM-III-170 are shown. The means and standard deviations derived from triplicate measurements within a typical experiment are shown. The experiments were repeated with comparable results.

In all four examples above, the presence of a single protomer with a PTC-destabilizing change apparently resulted in PTC destabilization of Env trimers with at least two protomers containing PTC-stabilizing changes. This is not surprising in the case where the PTC-stabilizing change is Q3, which even when present in all three protomers only weakly stabilizes the Env PTC. However, it is unexpected in the three instances with the strongly PTC-stabilizing Tri FPPR changes that, when mixed with wild-type HIV-1_AD8_ protomers, stabilize the PTC when present in a single protomer (N_p_=1) (Fig. 7B). Instead, when Tri FPPR protomers are mixed with Env protomers with PTC-destabilizing changes, Env trimers with one or two Tri FPPR protomers are cold sensitive. Apparently, the N_p_=1 stoichiometry of the PTC-destabilizing changes is dominant over the N_p_=1 stoichiometry of the PTC-stabilizing Tri FPPR change. The above examples also demonstrate that the Env background in which PTC-destabilizing changes are introduced may preserve or alter the protomer stoichiometry (N_p_).

### Sensitivity of Env chimeras to CD4mcs

The measurement of viral cold sensitivity affords a ligand-free means of evaluating the HIV-1 Env conformational state, thus providing insight into the <u>spontaneous</u> triggerability of Env. Virus sensitivity to CD4mc inhibition provides a second surrogate for the stability of the PTC, allowing an assessment of <u>induced</u> triggerability. Measuring the protomer stoichiometry of Env changes in the latter case is more technically challenging, as multiple CD4mc concentrations must be empirically evaluated to achieve detectable but different levels of infection for the wild-type and mutant viruses. We determined the protomer stoichiometry of three PTC-stabilizing Env changes (Q114E, Tri FPPR and FPPR-N136E/D325Q), using virus inhibition by the CD4mc BNM-III-170 as a phenotypic readout. None of these PTC-stabilizing changes overlaps with the known gp120 binding site of the CD4mcs (31–33). We mixed HIV-1_AD8_ Envs containing each of these changes with the wild-type HIV-1_AD8_ Env and tested the susceptibility of the viruses to inhibition by different concentrations of BNM-III-170. As expected (20,25), compared with viruses with the wild-type HIV-1_AD8_ Env, the viruses with 100% PTC-stabilized Envs were relatively resistant to BNM-III-170 inhibition. For the strongly PTC-stabilized Q114E and Tri FPPR Envs, the f-P curves exhibited mild shape changes as the BNM-III-170 concentration varied; these may reflect the effects of CD4mc occupancy of the gp120 subunits on the stoichiometric requirements for the PTC-stabilizing changes (Fig. 7A and B). However, the f-P curves for both Envs deviated only modestly from the N_p_=2 theoretical curve at all CD4mc concentrations. An N_p_=2 stoichiometry was also supported by the f-P curve observed for the PTC-stabilizing Tri FPPR changes when a more potent CD4mc, CJF-III-288 (33), was used (Fig. 7C). For the moderately PTC-stabilizing FPPR-N136E/D325Q Env, the f-P curves were consistent with N_p_=3 at multiple BNM-III-170 concentrations (Fig. 7D). Thus, for CD4mc resistance, we assign N_p_ values of 2, 2 and 3 for the Q114E, Tri FPPR and FPPR-N136E/D325Q Envs, respectively. The respective N_p_ values for the cold resistance of these mutants were 1, 1 and 2. Apparently, more protomers require PTC-stabilizing changes to achieve resistance to BNM-III-170 than to increase resistance to cold. It may be easier for PTC-stabilizing changes to resist spontaneous Env transitions from the PTC than the transitions induced by CD4mcs.

We also wished to evaluate how the presence of a PTC-destabilizing change in one or more Env protomers would affect virus sensitivity to BNM-III-170. For this purpose, we mixed HIV-1_AD8_ Envs containing the destabilizing N301E change with the wild-type HIV-1_AD8_ Env and tested the susceptibility of the viruses to inhibition by BNM-III-170. As expected (24), viruses with 100% AD8-N301E Envs were inhibited to a greater extent by BNM-III-170, consistent with their increased triggerability. At low concentrations of BNM-III-170, the viruses with two or more AD8-N301E protomers were inhibited (Fig. 7D). At higher BNM-III-170 concentrations, Envs with one protomer of the AD8-N301E Env were efficiently inhibited. Apparently, the CD4mc-induced inactivation of Env is more efficient when two protomers contain the PTC-destabilizing N301E change. Fewer Env protomers with this PTC-destabilizing change are required for virus inactivation when CD4mc concentrations are higher.

## DISCUSSION

In this work, we determined the number of Env protomers (N_p_) that must be modified by particular Env changes that stabilize or destabilize the pretriggered conformation (PTC) to achieve the viral phenotypes of increased resistance or sensitivity, respectively, to cold (0°C) or CD4mc exposure. Cold exposure and incubation with CD4mcs drive HIV-1 Env from the metastable PTC by different means: cold inactivation is ligand-free and spontaneous, whereas CD4mcs bind gp120 and induce changes in Env similar to those triggered by CD4 binding (11,12,19,23,26,33–36). Increases or decreases in the stability of the Env PTC result in increased HIV-1 resistance or sensitivity, respectively, to both cold and CD4mcs (12,17–26,34). The product of the virus half-life on ice and the CD4mc IC_50_ is the pretriggered Env stability index, which correlates with the stability of the PTC better than either factor on its own (25). We used the pretriggered Env stability indices to rank the HIV-1_AD8_ Env mutants in this study with respect to the degree of PTC stabilization or destabilization achieved.

The ligand-free cold inactivation assay allowed us to evaluate the cold sensitivity of viruses pseudotyped with mixtures of a great variety of Envs and to estimate the protomer stoichiometry. With respect to cold resistance of the viruses, the number of Env protomers that must be modified to achieve stabilization of the PTC is inversely correlated with the degree of PTC stabilization that results from the introduced Env changes (Fig. 4B). For strongly PTC-stabilizing Env changes, modification of a single protomer is sufficient to achieve PTC stabilization (Fig. 6). The latter observation suggests that given adequate stability, the PTC-stabilized protomer can influence the conformation of the other two protomers to maintain the PTC. The efficacy of such cross-protomer cooperativity is dependent on the Env partner that is mixed with the strongly PTC-stabilized Env. Thus, for the strongly PTC-stabilizing Tri FPPR changes, N_p_=1 when mixed with the wild-type HIV-1_AD8_ Env, but N_p_=3 when the Env partner contains PTC-destabilizing changes (N301E, Q3alt or S546D) (Fig. 7B). In the latter cases, the presence of even a single protomer with the PTC-destabilizing changes completely nullified the ability of the Tri FPPR changes to render the mixed Env trimers cold resistant. Apparently, disruption of the PTC of a single Env protomer is not readily compensated even by strongly PTC-stabilizing changes in the other protomers. In this situation, the protomer stoichiometries of PTC-destabilizing Env changes are dominant.

For weakly PTC-stabilizing Env changes, all three protomers required modification to achieve cold resistance of the virus (N_p_=3). Thus, symmetrical placement of weakly PTC-stabilizing changes in three Env protomers is apparently conducive to the maintenance of the PTC (Fig. 6). A complementary observation is that the presence of weakly PTC-destabilizing Env changes in one protomer was sufficient to increase the cold sensitivity of the virus (N_p_=1). The PTC of the wild-type HIV-1_AD8_ Env can be disrupted by the asymmetric introduction of a PTC-destabilizing Env change into a single protomer. This PTC-destabilizing N_p_=1 stoichiometry dominated, even when strongly PTC-stabilizing changes were present in the other protomers. In current Env structures (37–42), several of the strongly PTC-stabilizing changes involve amino acid residues that are near subunit interfaces. However, PTC-destabilizing changes often involve changes in N-linked glycosylation sites and are located on the periphery of the Env trimer (see below). Thus, direct interactions between Env structures involved in PTC stabilization and destabilization are unlikely explanations for our observations. A more attractive model explaining these observations is that the PTC represents a C3-symmetric trimer. In this model, Env changes that stabilize the PTC need to maintain conformational symmetry among the protomers. Weakly PTC-stabilizing Env changes, lacking the ability of the strongly PTC-stabilizing changes to act across protomers, must be present in all three protomers (N_p_=3). Conversely, PTC-destabilizing Env changes in one protomer can lead to a significant loss of conformational symmetry among the protomers, thereby destabilizing the PTC, with consequent increases in virus sensitivity to cold inactivation and CD4mc inhibition. Supporting this model are the many examples of weakly PTC-stabilizing Env changes where N_p_=3 and weakly PTC-destabilizing Env changes where N_p_=1 (Fig. 6).

Several of the weakly PTC-stabilizing changes (N136E, T138A, N136E/D325Q and Q3) involve the loss of a gp120 V1 glycan at Asn 136, which presumably exerts its effects at the Env surface (24,25). Other weakly PTC-stabilizing changes (Q567K and D325Q) do not alter a potential Env glycosylation site. Apparently, these changes most effectively stabilize the PTC when they are present in all three Env protomers, consistent with the stabilization of a symmetric Env trimer.

For the group of strongly PTC-stabilizing Env mutants (Q114E, Tri and Tri FPPR), all of which share the Q114E change, alteration of a single Env protomer is apparently sufficient to achieve cold resistance. In available Env trimer models (37–42), Gln 114 is located in the α1 helix of the gp120 inner domain, near the interface with gp41. The mechanism whereby a change in Gln 114 to glutamic acid in one protomer would stabilize the PTC of the entire Env trimer is not apparent in currently available structures. Detailed structures of the membrane Env PTC may shed light on this gap in our understanding.

For most of the PTC-destabilizing Env changes, alteration of a single Env protomer (N_p_=1) was apparently sufficient to sensitize the virus to cold inactivation. Loss of a glycan in one protomer (N_p_=1) destabilized the PTC in the case of the N301E and N332T changes, which remove glycans that occupy the interprotomer angles of the Env trimer (24). An N_p_ value of 1 was also deduced for the F317W change in the gp120 V3 tip, which does not alter an N-linked glycosylation site. The protomer stoichiometry of these Env variants supports the hypothesis that C3 symmetry contributes to the maintenance of the pretriggered (State-1) Env trimer conformation (Fig. 6). Recent structural studies suggest that the default intermediate (State-2) Env is an asymmetric trimer, which presumably derives from a symmetric pretriggered (State-1) Env (37,38,43).

One PTC-destabilizing change, S546D, needed to be present on two or more protomers (N_p_=2) to render the virus cold-sensitive. An obvious structural explanation is lacking for this requirement. Serine 546 is located in the highly dynamic gp41 HR1_N_ region, near the trimeric coiled coil formed by HR1_C_ (37–42). The higher N_p_ value of S546D, relative to those of other PTC-destabilizing changes, may reflect the magnitude of PTC destabilization achieved; the global sensitivity of the S546D mutant to antibody neutralization suggests that significant redistribution into State-3-like Envs results from this change.

To achieve HIV-1_AD8_ resistance to CD4mcs, the PTC-stabilizing changes were required in more Env protomers than for cold resistance. Apparently, it is more difficult to resist the direct induction by CD4mcs of entry-related transitions from the PTC than to withstand the more generally disruptive effects of ice formation on Env trimer integrity associated with cold exposure (35,36). With respect to CD4mc sensitivity, a PTC-destabilizing change, N301E, exhibited different protomer stoichiometries at low (N_p_=2) and high (N_p_=1) concentrations of the CD4mc. The increased occupancy of Env at higher CD4mc concentrations itself promotes Env transitions from the PTC and likely accounts for the lower protomer requirements for the PTC-destabilizing change.

Our studies of Env protomer stoichiometry will guide future efforts to evaluate the importance of Env trimer symmetry to PTC integrity and should facilitate the stabilization and characterization of the pretriggered (State-1) Env conformation.

## MATERIALS & METHODS

### HIV-1 Env mutants

The wild-type HIV-1_AD8_ *env* cloned in the pSVIIIenv expression plasmid was used as a template to construct HIV-1 Env mutants in this study (20). The signal peptide/N-terminus (residues 1-33) and the cytoplasmic tail C-terminus (residues 751-856) of this Env are derived from the HIV-1_HXBc2_ Env. “Tri” indicates the Q114E/Q567K/A582T changes, and “FPPR” indicates the A532V/I535M/L543Q changes. The Q3 and Q3 alt Env mutants were previously reported as Q3 (V1) and Q3 (V1alt), respectively, in reference 24. Env mutants with specific changes were generated by using the QuikChange Lightning site-directed mutagenesis kit (Agilent Technologies). All the Envs contain a His_6_ tag at the carboxyl terminus. The presence of the desired mutations was confirmed by DNA sequencing.

### Cell lines

HEK293T, TZM-bl and HOS cells (ATCC) were cultured in Dulbecco modified Eagle medium (DMEM) supplemented with 10% fetal bovine serum (FBS) and 100 mg/mL penicillin-streptomycin (Life Technologies).

### Env expression

To evaluate Env processing and subunit association, 3×10^5^ HOS cells were seeded in 6-well plates. After 24 hours of incubation, they were transfected with plasmids encoding His6-tagged Env variants and Tat at a ratio of 8:1, using the Lipofectamine 3000 transfection reagent (Life Technologies) according to the manufacturer’s instructions. Seventy-two hours after transfection, the cells were lysed in PBS buffer containing 1.0% NP-40 and protease inhibitor (Sigma-Aldrich). Clarified lysates were harvested, boiled, and analyzed by Western blotting with 1:2,500 goat anti-gp120 antibody (Invitrogen) and 1:2,500 HRP-conjugated rabbit anti-goat antibody (Invitrogen). The intensities of the gp120 and gp160 bands from unsaturated Western blots were quantified by using ImageJ software. The Env processing index was calculated by dividing gp120 by gp160 in the cell lysate samples. The processing indices of Env mutants in this study were normalized to those of the wild-type HIV-1_AD8_ Env.

Seventy-two hours after transfection, the supernatants of the transfected HOS cells were collected and incubated with Galanthus Nivalis Lectin (GNL)-agarose beads (Vector Laboratories) for 1.5 h at room temperature. The beads were washed three times with PBS containing 0.1% NP-40 and processed for Western blotting with 1:2500 goat anti-gp120 antibody (Invitrogen) as described above. The subunit association index was calculated by dividing gp120 in the cell lysate samples by the gp120 in the GNL precipitates. The subunit association indices of the Env mutants were normalized to those of the wild-type HIV-1_AD8_ Env.

### Virus infectivity

To produce pseudoviruses, HEK293T cells were cotransfected with the Env-expressing pSVIIIenv plasmid (a mixture of wild-type HIV-1_AD8_ Env with either PTC-stabilized or PTC-destabilized mutant Envs in varying proportions), a Tat-encoding plasmid and the luciferase-encoding pNL4-3.Luc.R-E-vector (NIH HIV Reagent Program) at a 1:1:3 ratio using polyethyleneimine (PEI, Polysciences). After 8 hours, the medium was replaced with fresh medium. Seventy-two hours later, the pseudoviruses in the supernatants were harvested and centrifuged (3500 rpm for 5 min), aliquoted, and either used directly to measure pseudovirus infectivity or stored at -80°C until further use. To evaluate the infectivity of the variants, pseudoviruses were diluted in 96-well plates and incubated with TZM-bl cells in the presence of 20 µg/mL DEAE-dextran. After 48 hours incubation, TZM-bl cells were lysed and the luciferase activity was measured using a luminometer.

### Virus sensitivity to cold inactivation

To evaluate the sensitivity of viruses with mixed Envs to cold inactivation, pseudoviruses were incubated on ice (0°C) for a period of time. This period of time was chosen to maximize the infectivity difference between viruses with wild-type HIV-1_AD8_ Env and PTC-stabilized/destabilized mutant Envs. After cold incubation, the infectivity of the pseudoviruses was measured as described above. The level of infectivity of each virus following incubation on ice was normalized to that of the same virus preparation that had not been incubated on ice.

### Virus inhibition by a CD4mc

**h**The CD4mcs BNM-III-170 and CJF-III-288 were serially diluted in triplicate wells in 96-well plates. Then approximately 100 to 200 TCID_50_ (50% tissue culture infectious dose) of pseudoviruses was added and incubated at 37°C for 1 h. Subsequently, approximately 2 × 10^4^ TZM-bl cells with 20 µg/mL DEAE-dextran in the medium were added to each well and the mixture was incubated at 37°C/5%CO_2_ for 48 hours. Then luciferase activity was measured, as described above. The level of infectivity of each virus with mixed Envs was normalized to that of the same virus preparation that had not been incubated with CD4mcs.

### Analysis of Env on virus particles

Approximately 1-mL of clarified cell supernatant containing pseudovirus was centrifuged at 14,000 × g for 1 h at 4°C. The pelleted virus particles were resuspended in 1x PBS. Equal volumes of the virus suspensions were then analyzed by Western blotting. Western blots were developed with 1:2,500 goat anti-gp120 polyclonal antibody (Invitrogen), 1:2,500 4E10 anti-gp41 antibody, and 1:5,000 rabbit polyclonal antibody against Gag p55/p24/p17 (Abcam). The HRP-conjugated secondary antibodies were 1:2,500 rabbit anti-goat antibody (Invitrogen), 1:2,500 goat anti-human antibody (Invitrogen), and 1:5,000 goat anti-rabbit antibody (Sigma-Aldrich), respectively.

## ACKNOWLEDGMENTS

We thank Ms. Elizabeth Carpelan for manuscript preparation. Antibodies against the HIV-1 Env were kindly supplied by Dennis Burton (Scripps), Peter Kwong and John Mascola (Vaccine Research Center, NIH), Susan Zolla-Pazner (Icahn School of Medicine at Mount Sinai), James Robinson (Tulane) and Hermann Katinger (Polymun). We thank the NIH HIV Reagent Program for providing additional reagents. This work was supported by grants from the National Institutes of Health (grant nos. AI145547, AI124982, AI150471, AI129017, AI164562, AI176904 and AI178833), a grant from Gilead Sciences, by Michael Siff Funds for Basic Research (Dana-Farber Cancer Institute) and by a gift from the late William F. McCarty-Cooper.

We declare no conflicts of interest.

